# Substrate elasticity does not impact on DNA methylation changes during differentiation of pluripotent stem cells

**DOI:** 10.1101/2024.01.16.575833

**Authors:** Mohamed H. Elsafi Mabrouk, Kira Zeevaert, Ann-Christine Henneke, Catharina Maaßen, Wolfgang Wagner

**Author notes:** Correspondence: Wolfgang Wagner. Those authors contributed equally to the work (Mohamed H. Elsafi Mabrouk, Kira Zeevaert).

## Abstract

Substrate elasticity may direct cell-fate decisions of stem cells. However, it is largely unclear how matrix stiffness impacts on differentiation of induced pluripotent stem cells (iPSCs) and if this is also reflected by epigenetic modifications. We have therefore cultured iPSCs on tissue culture plastic (TCP) and polydimethylsiloxane (PDMS) with different Young’s modulus (0.2 kPa, 16 kPa, or 64 kPa) to investigate the sequel on growth and differentiation towards endoderm, mesoderm, and ectoderm. Immunofluorescence and gene expression of canonical differentiation markers was hardly affected by the substrates. Notably, when we analyzed DNA methylation profiles of undifferentiated iPSCs or after three-lineage differentiation, we did not see any significant differences on the three different PDMS elasticities. Only when we compared DNA methylation profiles on PDMS-substrates *versus* TCP, we observed epigenetic differences, particularly upon mesodermal differentiation. Taken together, stiffness of PDMS-substrates did not impact on directed differentiation of iPSCs, whereas the moderate epigenetic differences on TCP might also be attributed to other chemical parameters.

## Introduction

Physical parameters - such as surface chemistry, micro-topography, or substrate elasticity - can impact on growth and differentiation of stem cells *in vitro* (1, 2). It is often anticipated that stem cells can sense the mechanical properties of their environment and differentiate accordingly. For example, it has been suggested that very soft tissues like the brain, with a Young‘s modulus of less than 1 kPa, direct toward neurogenic lineage; 8-17 kPa might support myogenic; and 25-40 kPa might rather correspond to cross-linked collagen of bones and hence enhance osteogenic differentiation (3). The effects of substrate elasticity were extensively investigated for adult stem cells, such as mesenchymal stromal cells (MSCs) (3, 4), whereas effects on pluripotent stem cells were less studied. For the latter, it has been suggested that substrate elasticity seems to alter the expression of germ layer markers during directed differentiation (5, 6) and affect differentiation of iPSC-derived embryoid bodies (7). On the other hand, there are studies that did not observe any effect of elasticity on directed germ layer specification of iPSCs (8, 9). Moreover, descriptions of unequivocal effects of substrate elasticity on differentiation of pluripotent cells is hampered by different substrate chemistries, topography, or 2D *versus* 3D approaches (5, 7, 10, 11). Thus, it is still under debate if matrix elasticity really is a central parameter to direct differentiation of pluripotent cells – and if potential phenotypic modifications would also be reflected in the epigenetic makeup, which ultimately defines cellular differentiation (9, 12). DNA methylation (DNAm) is an epigenetic modification that changes in a highly concerted and reproducible manner during cellular differentiation (13). In fact, the DNA methylation pattern is very cell type specific (14, 15), and it can be used to track early differentiation events of iPSCs (16, 17). We have therefore systematically investigated direct differentiation of iPSCs toward specific germ layers on polydimethylsiloxane (PDMS) with different Young’s modulus (0.2 kPa, 16 kPa, or 64 kPa), along with TCP, as a control material. If the elasticity of PDMS directs cell fate decisions, we would anticipate finding reproducible changes in the cell-type specific DNA methylation patterns.

## Materials and Methods

### Cell culture and differentiation of induced pluripotent stem cells

Three human induced pluripotent stem cell (hiPSCs) lines were used in this study with hPSCreg accessions (UKAi009-A, UKAi010-A, and UKAi011-A). They were generated from bone-marrow derived MSCs (iPSC-102-2, 104-12, and 106-3) and reprogrammed using episomal plasmids expressing *OCT4, L-Myc, SOX2, KLF4*, and *Lin28A* (9). The study was approved by the local ethic committee and all samples were taken after written consent (permit number: EK206/09). Cells were cultured regularly on tissue culture plastic (TCP) coated with vitronectin (0.5 µg/cm2) in StemMACS iPS-Brew (Miltenyi Biotec, Bergish Gladbach, Germany) and passaged regularly using intermittent Ethylenediaminetetraacetic acid (EDTA) treatment.

To investigate the impact of substrate elasticity, iPSCs were dissociated with Accutase (STEMCELL Technologies, Vancouver, Canada) and seeded (0.5-3×105 cells/cm2) on either PDMS with different Young’s modulus (0.2 kPa, 16 kPa, or 64 kPa; Advanced Biomatrix, Carlsbad, CA, USA) or TCP (approximately 1 GPa; TPP, Trasadingen, Switzerland), with Matrigel coating. Differentiation towards endoderm, mesoderm was carried out for 5 days and ectoderm for 7 days using STEMdiff Trilineage Differentiation Kit (STEMCELL Technologies). Alternatively, endoderm differentiation was performed according to the protocol by Wang et al. (18) on vitronectin-coated substrates. The endoderm differentiation medium consisted of RPMI-1640 base medium (Gibco, Carlsbad, USA) supplemented with 1% non-essential amino acids, L-Glutamine, and Penicillin/Streptomycin (all from Gibco) with addition of 1% B27 supplements (Gibco), CHIR99021 (6.45 µM; Tocris Biosciences, Bristol, UK), and Activin A (100 ng/mL; Proteintech, Rosemont, USA) for the first day, and only Activin A (100 ng/mL) for the following three days.

### Immunostaining and image analysis

Cells were fixed using 4% paraformaldehyde (PFA) for 10 minutes, treated with 0.1% Triton X-1000 (Carl Roth GmbH, Karlsruhe, Germany) and 1% bovine serum albumin (BSA; Sigma-Aldrich, Saint-Louis, USA) in PBS for 20 minutes, and then incubated overnight at 4°C with primary antibodies against OCT4 (clone C-10), PAX6 (clone AD2.35; both from Santa Cruz), Brachyury (R&D Systems), or GATA6 (clone D61E4; Cell Signaling). Secondary antibody staining was carried out at room temperature for 1 hour using donkey anti-goat (Alexa Fluor 488), goat anti-rabbit (Alexa Fluor 594), and goat anti-mouse (Alexa Fluor 594; all from Invitrogen). Samples were counterstained with DAPI for 15 minutes. Imaging was performed using the Axioplan 2 fluorescence microscope (Zeiss, Oberkochen, Germany) or EVOS FL Auto (Life Technologies, Carlsbad, USA). Quantification of fluorescence images was carried out using cellpose python package (v2.2) (19). Fluorescence markers and DAPI images were segmented using the pre-trained “nuclei” model with default parameters and the number of cells was counted from the generated segmentation mask using scikit-image (20).

### Semi-quantitative reverse-transcription PCR

Total RNA was isolated using the NucleoSpin RNA kit (Macherey-Nagel, Düren, Germany), quantified with the NanoDrop ND-1000 spectrophotometer (Thermo Scientific, Waltham, USA), and reverse transcribed using the High-Capacity cDNA Reverse Transcription Kit (Applied Biosystems, Waltham, USA). Semi-quantitative Reverse Transcription PCR (RT-qPCR) was carried out using Power SYBR Green PCR Master Mix (Applied Biosystems, Waltham, USA) and primers for germ layer differentiation and pluripotency marker genes (Suppl. Table 1) using StepOnePlus machine (Applied Biosystems, Waltham, USA). *GAPDH* was used for normalization.

### DNA methylation analysis

Genomic DNA was isolated using the NucleoSpin Tissue Kit (Macherey-Nagel) and quantified on a NanoDrop ND-2000 spectrophotometer (Thermo Scientific). 1.2 µg of DNA were bisulfite converted and analyzed using the EPIC BeadChip microarray (Illumina, San Diego, USA) by Life & Brain company (Bonn, Germany). The data was analyzed with the minfi R package (21) and normalized using ssNoob (22). CpG sites on XY chromosomes, non-CG probes, and SNP associated CpGs were removed. In addition, the detection p-value of the remaining CpGs were evaluated using SeSAMe package and CpG sites with detection *P* > 0.05 were removed (23). Limma R package was used for identification of differentially methylated CpGs. Significant CpG sites were defined as showing at least 10% difference in mean beta values and a Benjamini-Hochberg adjusted p-value smaller to 0.05. Over-representation analysis of the significant CpGs was carried out using the MissMethyl package employing the gene ontology database (24). Estimation of germ layer specification was carried with GermLayerTracker (alternatively called PluripotencyScreen), which is based on DNA methylation at 12 CpGs (17). As SeSAMe masking is stringent and resulted in removal of several CpGs associated with the GermLayerTracker, we carried out the deconvolution using the more permissive detection p-values from the minfi package instead.

### Statistical analysis

Statistical analysis was carried out using t-test from statannotations python package for quantification of immunofluorescence imaging or the moderated t-test from the limma R package for differential methylation analysis. *P* was adjusted for multiple testing using the Benjamini-Hochberg procedure when appropriate and *P* > 0.05 were considered significant.

## Results

### Substrate elasticity hardly affects differentiation of iPSCs

Human iPSCs were cultured on soft (0.2 kPa), medium (16 kPa), or stiffer PDMS-substrates (64 kPa), and for comparison on conventional tissue culture plastic (about 1 GPa). Immunophenotypic analysis revealed similar expression of OCT4 (Fig. 1A), and absence or very low expression of the germ layer specific markers GATA6 (endoderm), Brachyury (mesoderm), and PAX6 (ectoderm, Suppl. Fig. S1A). When we induced differentiation towards endoderm, mesoderm, and ectoderm, we observed the clear presence of GATA6, Brachyury, and PAX6 protein, respectively – and there were again no clear differences on the different substrates (Fig. 1A). Furthermore, gene expression of pluripotency markers *OCT4* and *NANOG*; mesoderm markers *TBXT* (Brachyury) and *MIXL1;* and ectoderm markers *PAX6* and *LHX5* were similar on the different substrates (Fig. 1B, Suppl. Fig. S1B). Notably, a moderate up-regulation of the endodermal differentiation markers *GATA6* and *SOX17* was observed on 0.2 kPa substrates (n = 6; *P* < 0.05) as compared to differentiation on PDMS substrates of higher elasticity and TCP (Fig 1B). However, this could not be validated in immunofluorescence imaging, even after quantification of GATA6 positive cells by image segmentation using a neural network model (3 biological replicas; n > 20 per substrate; t-test; Suppl. Fig 1C).

**Figure 1:**
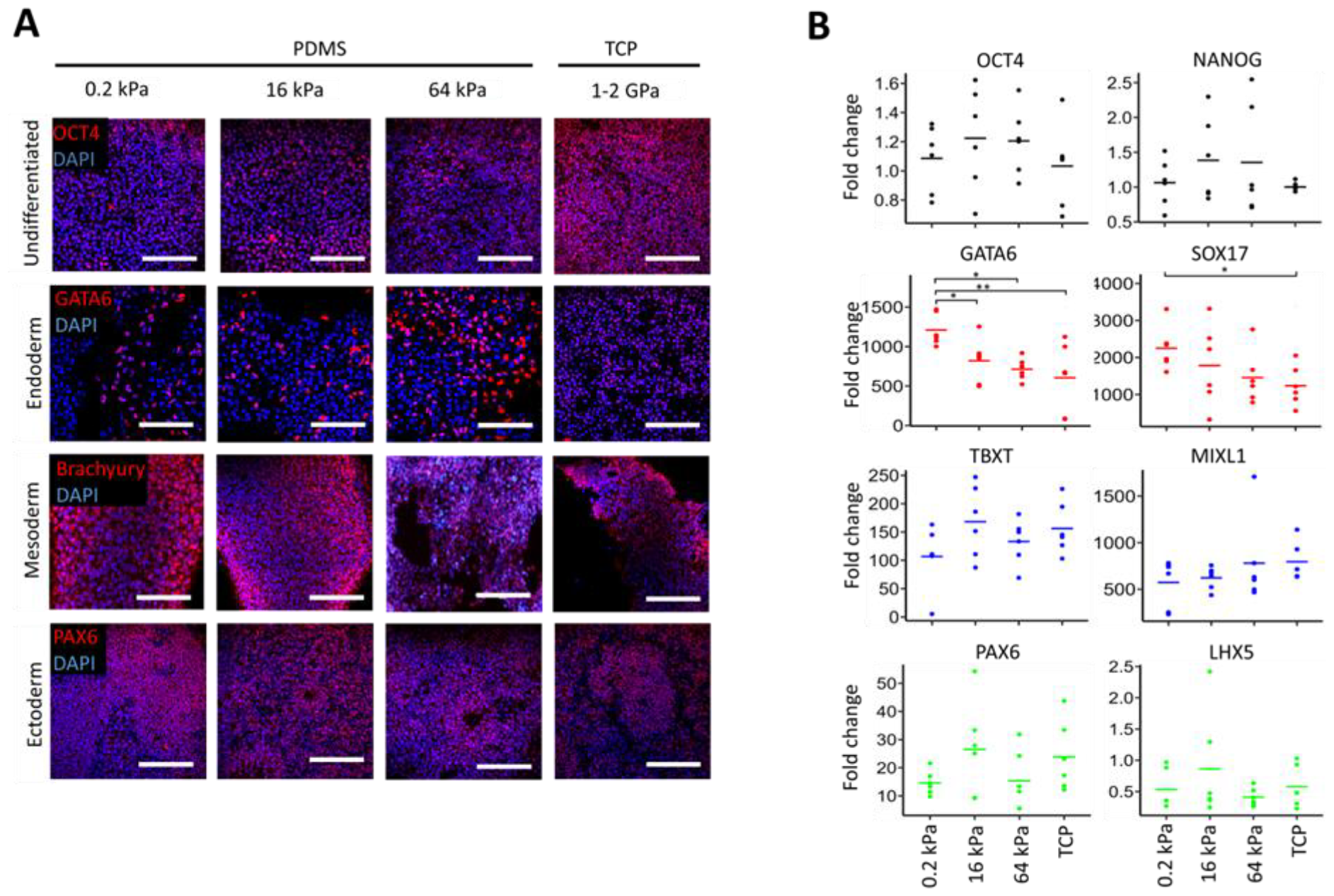
Early germ layer specification of iPSCs on different substrates. **A)** Immunofluorescence images of cells in PDMS substrates with different elasticities (0.2 kPa, 16 kPa, and 64 kPa), or tissue culture plastic (TCP). Undifferentiated iPSCs stained with OCT4, endoderm differentiation stained with GATA6, mesoderm differentiation stained with Brachyury, and ectoderm differentiation stained with PAX6. Nuclei were counterstained with DAPI. Images show no significant differences between undifferentiated or differentiated iPSCs on different substrate elasticities (scale bar: 50 µm). **B)** Gene expression analysis with RT-qPCR of candidate genes for lineage-specific differentiation. Data are normalized to undifferentiated cells on TCP (n = 6; significance measured with unpaired t-test; *p < 0.05, **p < 0.01).

We alternatively tested a different differentiation regimen toward endoderm on TCP and 64 kPa PDMS, and there was no significant difference in immunofluorescence of GATA6, nor did the coating with either vitronectin or Matrigel impact on the differentiation (n > 10; Suppl. Fig. S2A-B). The up-regulation of gene expression of *GATA6, SOX17*, or *NANOG* was also the same on PDMS and TCP (n = 6; Suppl. Fig. S2C). Taken together, they retained their stemness and revealed very similar differentiation on the different substrates, which is in line with previous observations (8).

### DNA methylation changes during differentiation are not affected by elasticity

Subsequently, we investigated how the DNA methylation profiles were affected by culture and differentiation of iPSCs on the different substrates. Multidimensional scaling (MDS) plot for the top 10,000 variable CpG sites revealed that samples mainly separated according to their state of differentiation (Fig 2A). Initially we focused on these germ layer associated epigenetic modifications across all different substrates and observed significant DNA methylation changes at 16,024 CpGs during endodermal differentiation, in 63,729 CpGs during mesodermal differentiation, and in 22,059 CpGs during ectodermal differentiation (delta mean DNAm >10%, Adj. *P* < 0.05; Fig. 2B). These epigenetic modifications were enriched in Gene Ontology (GO) categories that may be relevant for the corresponding differentiation process (Suppl. Fig. S3). Furthermore, GermLayerTracker analysis (17) could clearly classify all cell preparations as undifferentiated, endodermal, mesodermal and ectodermal differentiation – but, there was no clear impact from the substrates (Fig 2C).

**Figure 2:**
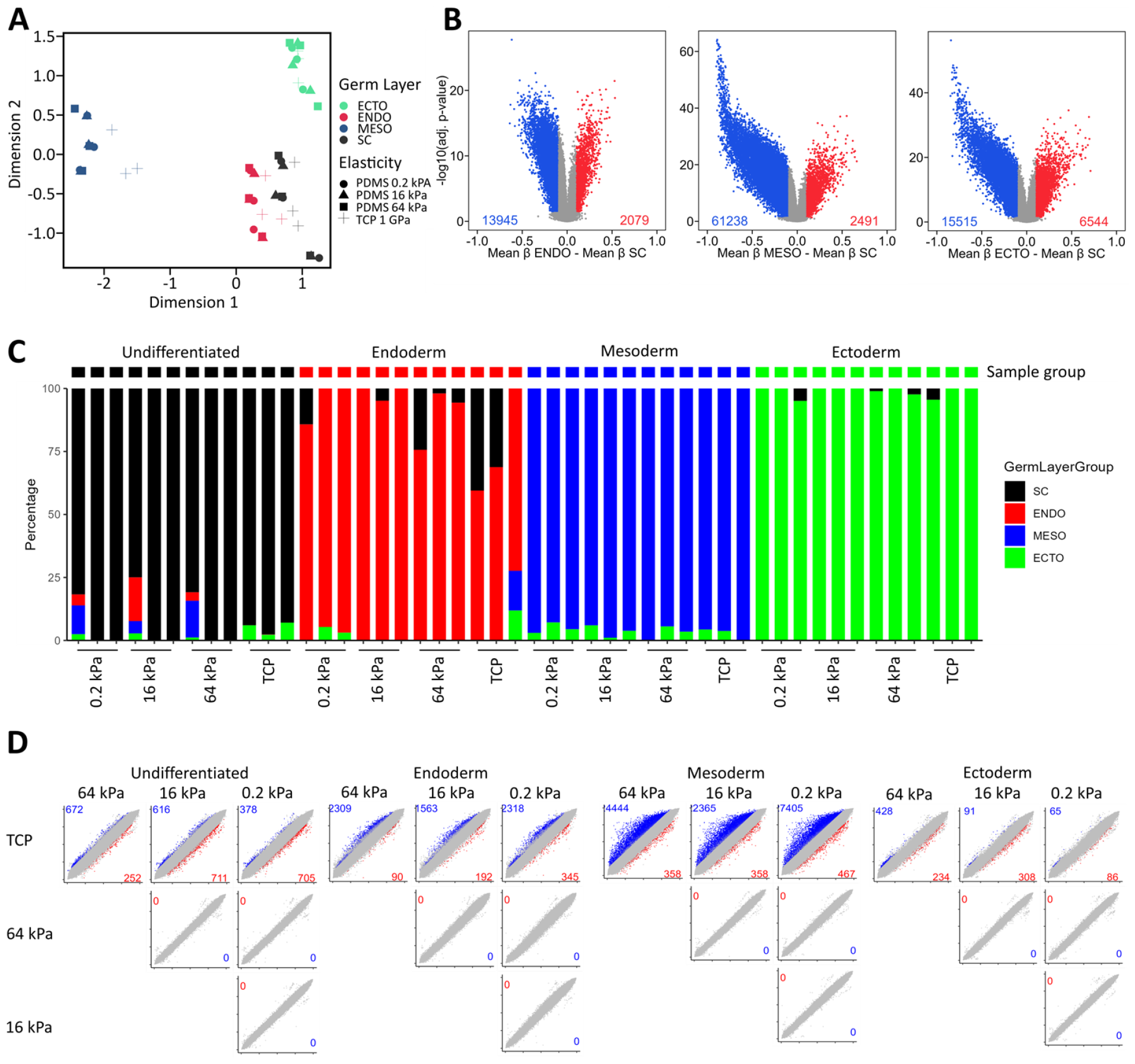
DNA methylation changes during germ layer specification on different elasticities. **A)** MDS dimensional reduction plot for top 10,000 variable CpG sites for ectoderm (ECTO), endoderm (ENDO), mesoderm (MESO), and iPSCs (SC). Cells were cultured on substrates with different Young’s modulus on PDMS (0.2 kPa, 16 kPa, and 64 kPa) and TCP (1 GPa). **B)** Volcano plot showing significantly hypermethylated (red) or hypomethylated (blue) CpGs after differentiation into endoderm, mesoderm, and ectoderm, respectively (Adj. p < 0.05; diff. in mean DNAm (beta-value) as compared to iPSCs > 10%). **C)** Estimation of the composition of germ layers based on epigenetic signatures of 12 CpGs of GermLayerTracker (PluripotencyScreen) (17). **D)** Pairwise comparison of differentiated CpGs between cells culture on different substrates. This analysis was performed for undifferentiated iPSCs, endoderm, mesoderm, and ectoderm differentiation. The scatter plots depict mean DNAm values on the corresponding substrates and the number of significantly hypo- and hypermethylated CpGs are indicated (Adj. p < 0.05, diff. mean DNAm > 10%).

To further investigate if substrate elasticity had significant impact on DNA methylation profiles, we performed pairwise comparison between the three replicates of each culture condition. There were some differences between TCP and each of the PDMS substrates. However, there were no significant differences upon culture on the different PDMS elasticities (Fig. 2D). This was unexpected as PDMS substrate covered a broad range of elasticities (from 0.2 kPa to 64 kPa). Thus, the different substrate elasticities in PDMS did neither affect DNA methylation patterns in undifferentiated iPSCs, nor the DNA methylation changes during directed differentiation.

### Different materials can evoke DNA methylation changes

To better understand the above-mentioned differences between PDMS and TCP, we compared all PDMS substrates *versus* TCP. Even under non-differentiation culture conditions, we observed 522 hypomethylated, and 514 hypermethylated CpGs on PDMS (Adj. p-value < 0.05; mean DNAm difference > 10%). Particularly upon mesodermal differentiation there were marked epigenetic differences between PDMS and TCP: 25,254 CpGs became significantly hypomethylated (Fig 3A). We then analyzed if these substrate-associated DNA methylation changes were related to the germ layer specific DNA methylation changes. In fact, the vast majority of the CpGs that were hypomethylated on PDMS were also hypomethylated during the course of mesodermal differentiation (Fig 3B). Thus, the choice of PDMS and TCP affects the DNA methylation profiles and may point toward enhanced mesodermal differentiation on PDMS.

**Figure 3:**
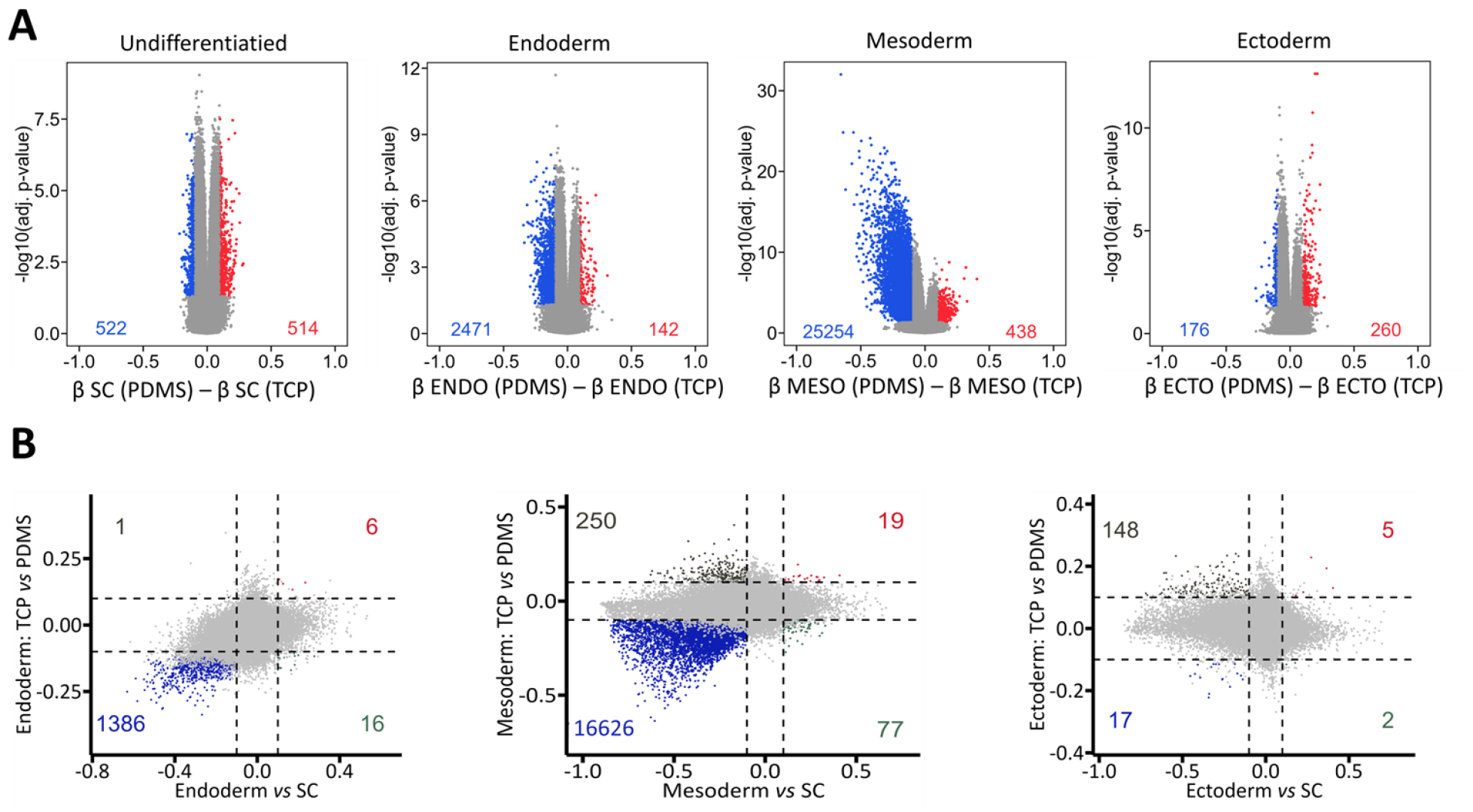
Differentially methylated CpGs on PDMS versus TCP. **A)** Volcano plots showing differentially methylated CpGs between samples cultured on TCP or PDMS for undifferentiated iPSCs, endodermal-, mesodermal-, and ectodermal-differentiated cells. The numbers of significantly hypomethylated (blue) and hypermethylated (red) CpGs on PDMS are indicated (Adj. p < 0.05, diff. mean DNAm (beta-value) > 10%). **B)** Scatter plots showing comparison of DNA methylation changes during differentiation toward endoderm, mesoderm, and ectoderm, respectively (x-axis), against the DNA methylation changes between TCP and PDMS (y-axis). Many CpGs that are hypomethylated during mesodermal differentiation are also hypomethylated on PDMS.

## Discussion

Within the last 15 years, it became almost a dogma that stiffness plays a central role in directing cell-fate decisions. Thus, it was unexpected that the broad range of substrate elasticities on PDMS did neither clearly affect differentiation of iPSCs, nor their DNA methylation patterns. It is conceivable, that some of the matrix stiffness related effects are masked by the interaction with proteins that are absorbed to the surface for coating or secreted by the stem cells (25, 26). Yet, culture of iPSC is not possible without suitable coating (27), and we did not observe differences on Matrigel or vitronectin coated substrates. Furthermore, in our previous work we did not observe significant epigenetic differences in iPSC-derived mesenchymal stromal cells (iMSCs) that were either generated on tissue culture plastic or a collagen-based hydrogel (9). It is plausible that some of the phenotypic stiffness-related effects are only transient – while the cells are on the substrate – but not reflected in epigenetic changes, which ultimately determine cell fate.

As already mentioned above, modulation of the physical parameter elasticity is often interwoven with other relevant factors, such as surface topography or surface chemistry (28, 29). This might explain, why we observed some epigenetic differences on TCP *versus* PDMS. Cells committed to mesoderm and endoderm were shown to undergo widespread coordinated epigenetic rearrangements driven by ten-eleven translocation (TET)-mediated demethylation (30). It needs to be further explored, if the increased hypomethylation events on PDMS substrates are really associated with enhanced mesodermal differentiation – at least with regards to immunophenotype or expression of marker genes, we did not observe such differences.

Taken together, a wide range of elasticities of PDMS substrates did not affect differentiation of iPSC. Albeit TCP has a very high non-physiological Young’s modulus, epigenetic impact of TCP *versus* PDMS is not necessarily related to differences in elasticity – they may rather be attributed to many other physical or chemical parameters that differ between these materials.

## Glossary

(DNAm): DNA methylation
(iMSCs): iPSC-derived mesenchymal stromal cells
(iPSCs): induced pluripotent stem cells
(MSCs): mesenchymal stromal cells
(PDMS): polydimethylsiloxane
(RT-qPCR): Semi-quantitative Reverse Transcription PCR
(TCP): tissue culture plastic
(TET): ten-eleven translocation
(MDS): Multi dimensional scaling

## Author Contributions

M.E.M. contributed to experimental procedure, performed image quantification and bioinformatics and data analysis. K.Z. performed iPSCs cell culture and differentiation experiments. A.C.H and C.M. contributed to gene expression measurement and immunofluorescence imaging. W.W. conceptualized and supervised the study. M.E.M. and W.W. wrote the manuscript and all authors approved final version.

## Supplemental Materials

### Supplemental Figures S1-S4 and Table S1

Fig S1: Early germ layer marker gene expression in undifferentiated iPSCs; Fig S2: Endoderm differentiation on substrates with different coating; Fig S3: Gene ontology for CpGs associated with germ layer differentiation; Fig. S4: Differential DNA methylation during germ layer differentiation versus PDMS/TCP; Table S1: Primer list.

