## Supplemental Figures S1-S4 and Table S1 for "Substrate elasticity does not impact on DNA methylation changes during differentiation of pluripotent stem cells"

1. Helmholtz-Institute for Biomedical Engineering, RWTH Aachen University Medical School, Aachen, 52074, Germany.
2. Institute for Stem Cell Biology, University Hospital of RWTH Aachen, 52074 Aachen, Germany.
* Those authors contributed equally to the work.


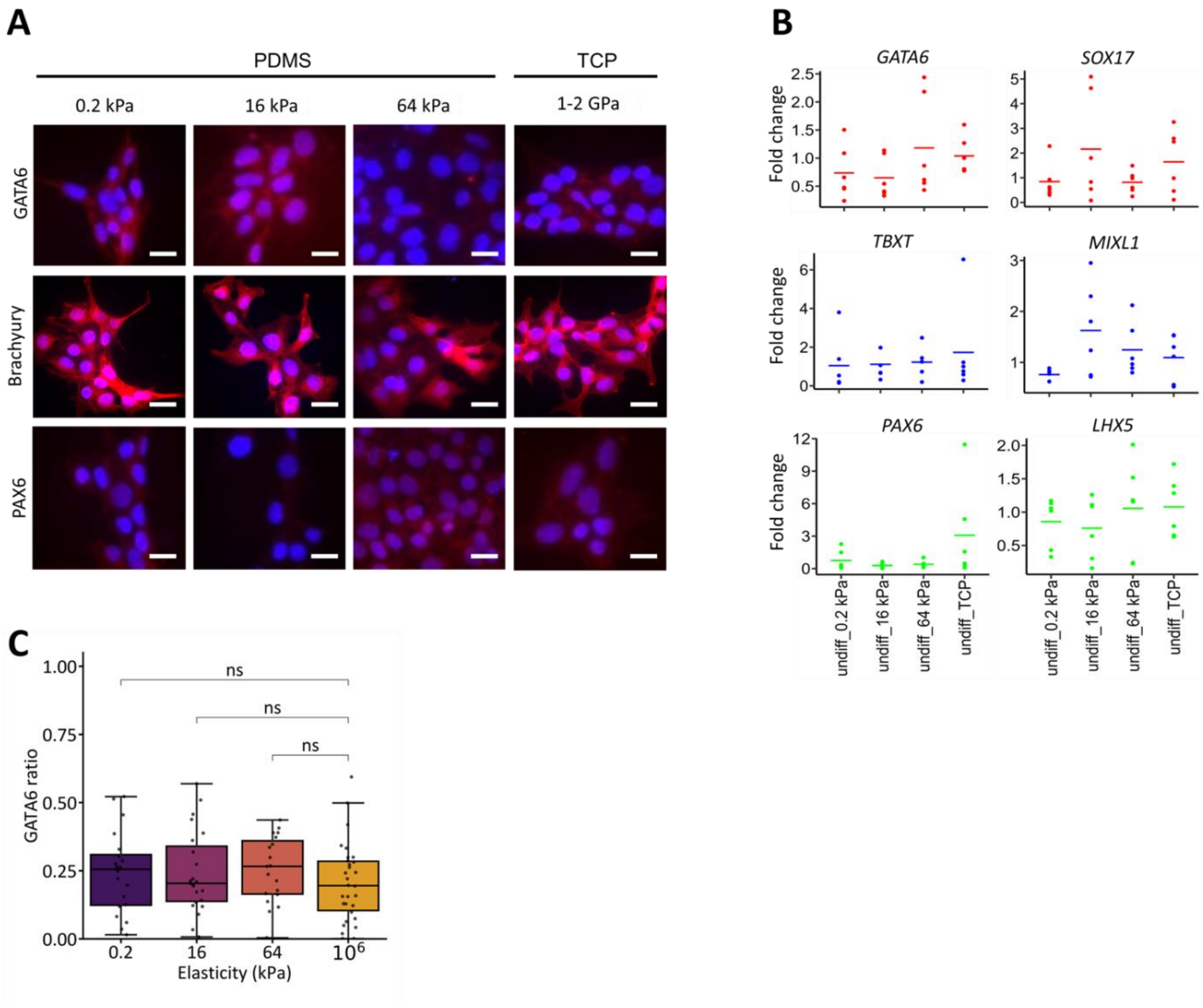


**Suppl. Fig. S1: Early germ layer marker gene expression in undifferentiated iPSCs.**
**A)** Immunofluorescence imaging of undifferentiated iPSCs cultured for 5 days on PDMS substrates with 0.2 kPa, 16 kPa, and 64 kPa elasticity; and TCP substrate (approximately 1 GPa). Samples are stained with antibodies against endodermal marker (GATA6), mesodermal marker (Brachyury), and ectodermal marker (PAX6). The images show no bias in the undifferentiated iPSCs towards specific germ layer based on substrate elasticity (scalebar: 50 µm). **B)** RT-qPCR for undifferentiated iPSCs analogous to A showing no significant differences in gene expression of germ layer markers based on substrate elasticity. **C)** Quantification of GATA6 immunofluorescence imaging using nuclear segmentation neural network model. GATA6 ratio is the fraction of nuclei stained with DAPI expressing GATA6 (n > 20; three biological replicas; significant measured using t-test; ns: not significant).

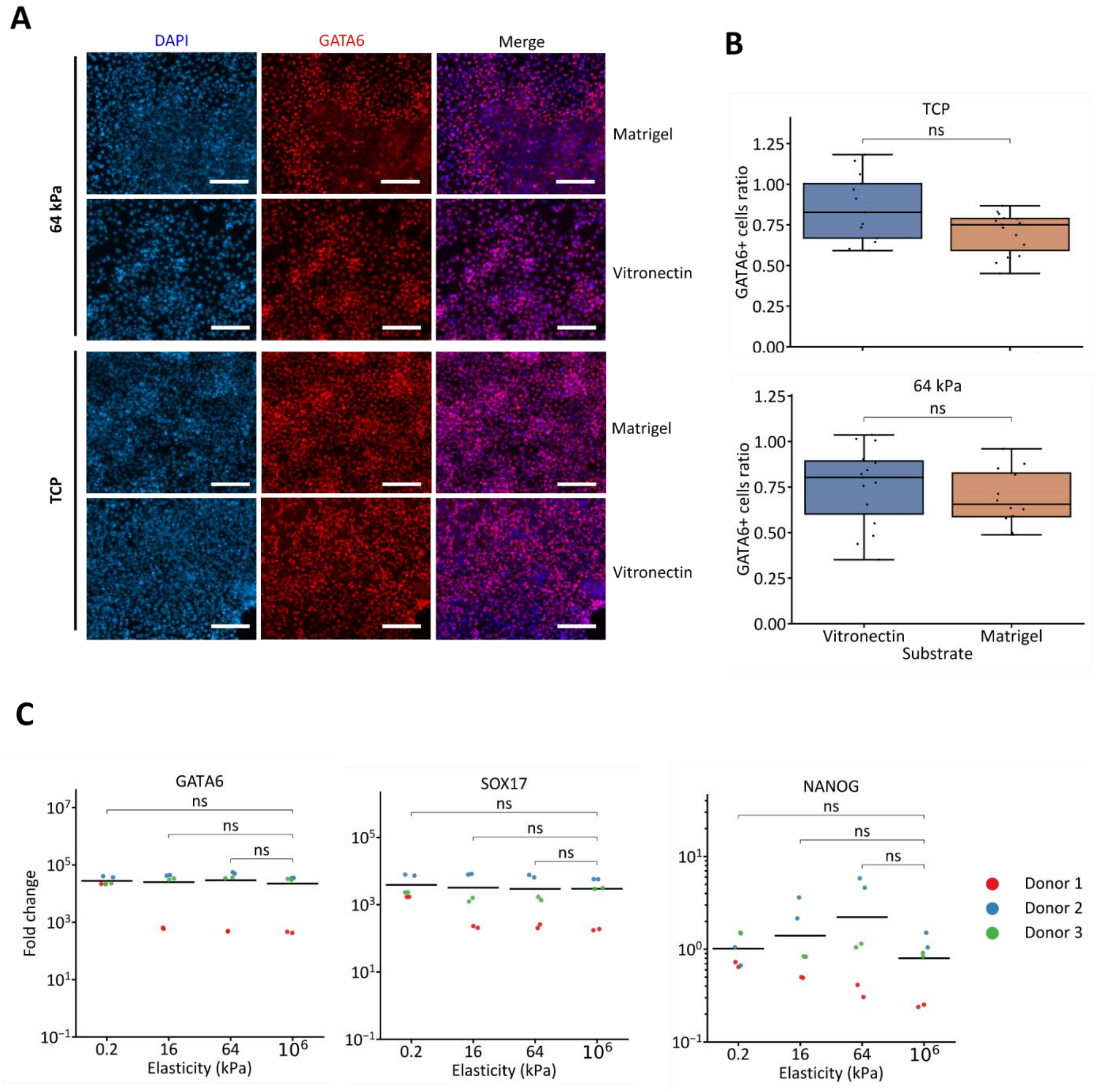


**Suppl. Fig. S2: Endoderm differentiation on substrates with different coating.**

**A)** Comparison of immunofluorescence imaging of endoderm marker GATA6 for iPSCs differentiated on vitronectin- or Matrigel-coated substrates with substrate elasticity of 64 kPa or TCP (scale bar: 200 µm). **B)** Quantification of immunofluorescence imaging of GATA6 using nuclear segmentation neural network model (n > 10; t-test; ns: not significant). **C)** RT-qPCR of endoderm marker genes *GATA6* and *SOX17*, and the pluripotency marker *NANOG* for endoderm differentiated cells on vitronectin-coated substrates of elasticities ranging from 0.2 kPa to TCP (1-2 GPa). (n = 6; three biological replicas; significance measured using t-test; ns: not significant).

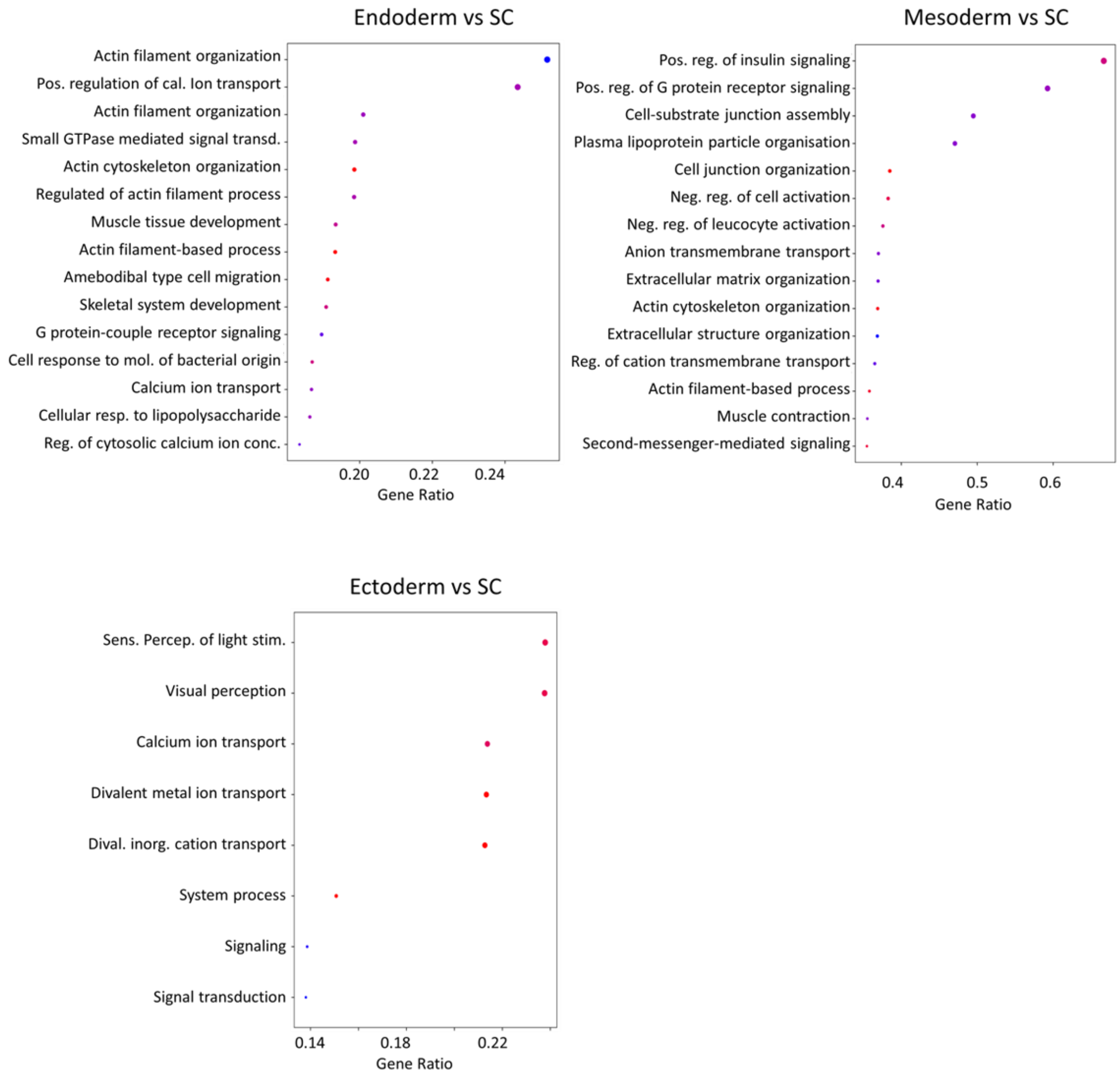


**Suppl. Fig. S3: Gene ontology for CpGs associated with germ layer differentiation.**

Gene ontology for CpGs associated with germ layer differentiation (as indicated in Fig. 2B) using MissMethyl package. Overall, DNA methylation changes are enriched in gene categories that may be involved in the corresponding differentiation processes ($p < 0.05$).

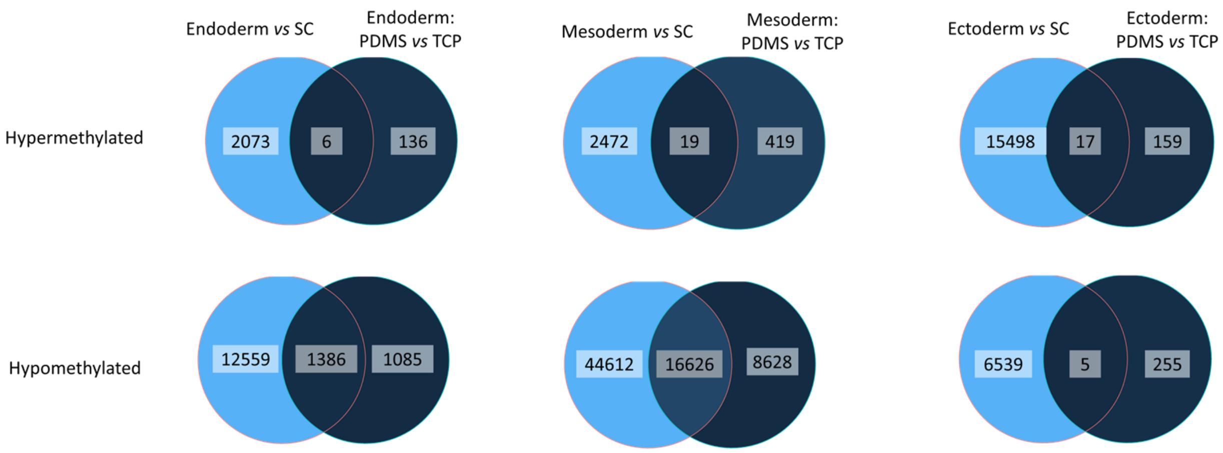


**Suppl. Fig. S4: Differential DNA methylation during germ layer differentiation *versus* PDMS/TCP.**
**A)** Venn diagrams showing shared differentially methylated CpGs between differentiation on TCP *versus* PDMS and iPSCs *versus* endodermal-, mesodermal-, and ectodermal differentiation. Mesoderm shows the highest number of shared CpGs with 16,626 shared hypomethylated CpGs between the two conditions.

**Suppl. Table S1: Primer list.**

| Target Gene | Full Name | Primer | DNA sequence |
|---|---|---|---|
| *POU5F1 (OCT4)* | POU Class 5 Homeobox 1 | Forward | GGGGGTTCTATTTGGGAAGGTA |
| | | Reverse | ACCCACTTCTGCAGCAAGGG |
| *NANOG* | Nanog Homeobox | Forward | CAGAAGGCCTCAGCACCTAC |
| | | Reverse | ATTGTTCCAGGTCTGGTTGC |
| *PAX6* | Paired Box 6 | Forward | TCGAAGGGCCAAATGGAGAAGAGAAG |
| | | Reverse | GGTGGGTTGTGGAATTGGTTGGTAGA |
| *LHX5* | LIM Homeobox 5 | Forward | GCGAGTGCAAAACCAACCTCT |
| | | Reverse | GGCGCATTTCGTGCCAAAG |
| *MIXL1* | Mix Paired-Like Homeobox | Forward | CGACATCCACTTGCGCGAG |
| | | Reverse | GGAAGGATTTCCCACTCTGACG |
| *TBXT* | T-Box Transcription Factor T | Forward | CAGTGGCAGTCTCAGGTTAAGAAGGA |
| | | Reverse | CGCTACTGCAGGTGTGAGCAA |
| *SOX17* | SRY-Box Transcription Factor 17 | Forward | AGGAAATCCTCAGACTCCTGGGTT |
| | | Reverse | CCCAAACTGTTCAAGTGGCAGACA |
| *GATA6* | GATA Binding Protein 6 | Forward | CTCAGTTCCTACGCTTCGCAT |
| | | Reverse | GTCGAGGTCAGTGAACAGCA |
| *GAPDH* | Glyceraldehyde-3-Phosphate Dehydrogenase | Forward | GAAGGTGAAGGTCGGAGTC |
| | | Reverse | GAAGATGGTGATGGGATTTC |
